# *CHD8* regulates the balance between proliferation and differentiation of human iPSCs in neural development

**DOI:** 10.1101/732693

**Authors:** Wenzhong Liu, Weilai Dong, Ellen J. Hoffman, Thomas V. Fernandez, Abha R. Gupta

## Abstract

**Background:** *Chromodomain helicase DNA-binding protein 8 (CHD8)*, which encodes a chromatin remodeling protein that regulates Wnt/β-catenin mediated gene expression, is one of the most strongly associated genes with autism spectrum disorder (ASD). Characterization of ASD patients with CHD8 disrupting mutations and animal and stem cell models of CHD8 deficiency suggest that CHD8 plays a role in neurodevelopment.

**Methods:** We generated iPSCs from the T-lymphocytes of a healthy, typically-developing human male and iPSC lines from the same source which were subjected to CRISPR/Cas9-mediated knockdown (KD) of *CHD8*. We subsequently derived neural progenitor cells (NPCs) and neural cells and examined the effects of CHD8 deficiency on cell proliferation and neural differentiation.

**Results:** We observed that, compared to WT, *CHD8* KD: (1) increased the number of iPSC colonies formed, (2) suppressed spontaneous differentiation along the edges of iPSC colonies, (3) increased the proliferation of NPCs, (4) delayed the formation of neural rosettes, (5) delayed neurite outgrowth, (6) decreased the percentage of cells in the G0/G1 phase of the cell cycle, (7) increased the percentage of cells in the G2/M phase of the cell cycle, (8) decreased presence of the neuronal marker MAP2 although not the glial marker GFAP, (9) decreased presence for the excitatory neuronal marker VGLUT1, and (10) decreased presence of the synaptic marker SYN1.

**Conclusions:** Our results suggest that CHD8 deficiency causes alterations in the cell cycle. More specifically, CHD8 KD appears to increase cell proliferation and delay neural differentiation. This may contribute to the pathophysiology of ASD.

## BACKGROUND

Autism spectrum disorder (ASD) is defined by deficits in social communication and interaction and restricted, repetitive patterns of behavior, interests, or activities (1). High-throughput sequencing studies have revealed that *de novo* sequencing and copy number variants make an important contribution to the genetic architecture of ASD (2-4). The identification of two dozen *de novo* mutations in *chromodomain helicase DNA-binding protein 8 (CHD8)* makes this gene one of the most strongly associated with ASD (5).

*CHD8* encodes an ATP-dependent chromatin remodeling protein that regulates Wnt/β-catenin mediated gene expression (6, 7). Indeed, transcriptional regulation has emerged as an important molecular pathway in the pathophysiology of ASD (2). Its expression peaks in the early prenatal period of human brain development but continues to be widely expressed throughout the adult brain (8).

Clinical characterization of ASD patients with *CHD8* mutations has revealed some common features, such as macrocephaly, distinct facial features, and gastrointestinal (GI) disturbances, suggesting a distinct ASD subtype (8). A number of animal models, including zebrafish and mice, have been generated to investigate *CHD8* deficiency (8-14). Whereas homozygous *Chd8* knockout (KO) mice die *in utero* (10), heterozygous *Chd8* KO mice are viable but have an array of abnormalities that provide insights into the human phenotype. Depending on the specific model and studies done, germline KO mice have macrocephaly (10-13); decreased GI motility (10); craniofacial abnormalities (11); ASD-like behaviors such as altered social behavior, repetitive behaviors, and increased anxiety (10); and/or cognitive impairment (12). Gene expression analysis revealed widespread changes in gene expression, and gene set enrichment analysis suggested delays in fetal neurodevelopment (10). Mutant mice generated by *in utero* KD of *Chd8* using shRNAs resulted in defective neural progenitor cell (NPC) proliferation and differentiation (14), in contrast to the macrocephaly phenotype. Possible reasons for the difference include the different methodologies used to generate the mice and differing effects of *CHD8* deficiency in neuronal and non-neuronal cells (14, 15). The precise role of *CHD8* in cellular proliferation and differentiation needs further delineation.

In addition to animal models, a number of studies have characterized *CHD8*-deficient stem cells. *CHD8* KD in NPCs derived from human induced pluripotent stem cells (iPSCs) resulted in down-regulated genes which are involved in neurodevelopment and are enriched for ASD-associated genes (9). A separate study showed that ASD-associated genes are overrepresented among *CHD8* target genes in the developing brain and that *CHD8* KD in human neural stem cells causes dysregulation of these genes (16). *CHD8* deficiency has been found to alter the expression of both protein-coding and noncoding RNAs (17). Two reports examined the transcriptional network in NPCs and monolayer neurons (18) and in cerebral organoids (19) derived from CRISPR/Cas9-mediated heterozygous KO of *CHD8* in human iPSCs. Both studies showed that differentially expressed genes were enriched for roles in neurodevelopment and Wnt/β-catenin signaling. Among these genes were those associated with human brain volume or head size (18) and the noncoding RNA *DLX6-AS1*, which regulates transcription factors involved in GABAergic interneuron differentiation (19). A third study using CRISPR/Cas9 genome editing was not able to generate homozygous *CHD8* KO in human iPSCs, suggesting that lack of CHD8 is lethal (20). This study did not obtain any significant results by phenotyping heterozygous *CHD8* KO cells using RNAseq or patch-clamp and multi-electrode array recordings.

Our aim is to further investigate the effects of *CHD8* KD in human iPSCs, NPCs, and neural cells. We compared the proliferation and differentiation of lymphocyte-derived iPSCs from a healthy, typically-developing human male and iPSC lines from the same source which were subjected to CRISPR/Cas9-mediated KD of *CHD8*.

## METHODS

### Generation of iPSCs from wild-type human T-lymphocytes

This research was approved by the Yale University Institutional Review Board. After obtaining written informed consent, a blood sample was drawn from a healthy, typically-developing, 25 year-old male subject. Peripheral blood mononuclear cells (PBMCs) were isolated by centrifugation of heparinized blood over a Ficoll-Paque PLUS gradient (GE Healthcare Life Sciences). PBMCs were cultured at 37°C in 5% CO_2_ and 10% FBS RPMI with anti-CD3/CD28 monoclonal antibody (STEMCELL Technologies) and 50 mU/ml IL-2 (PeproTech). After five days, activated T-lymphocytes were collected and transduced using the CytoTune-iPS Sendai Reprogramming kit, which contains Sendai viral vector encoding the Yamanaka factors cMyc, Klf4, Oct3/4, and Sox2 at MOI of 3:1 (Thermo Fisher Scientific). After 24 hours, the medium was changed to RPMI1640 containing IL-2. On day #8, transduced cells were placed on mouse embryonic fibroblasts (MEFs) in human iPSC medium [DMEM/F12 containing 20% KnockOut Serum Replacement (Thermo Fisher Scientific), 1% GlutaMAX-I (Gibco), 1% nonessential amino acids (Gibco), 1X β-mercaptoethanol (Gibco), and 4ng/ml FGF (PeproTech)]. The media was changed every other day. iPSC colonies appeared by week #3 and were transferred to Matrigel (BD) coated 24-well plates in mTeSR1 medium (STEMCELL Technologies) (Fig. S1). The medium was changed every other day, and the cells were passed every 6-7 days using Dispase (STEMCELL Technologies).

### Quality Control

Pluripotency was tested by staining for alkaline phosphatase (ALP) using ALP Live Stain (Life Technologies) and immunofluorescent staining for Tra-1-60, SSEA3, and SSEA4 (Fig. S2). RT-PCR was conducted on iPSC RNA to detect expression of cMyc, Klf4, NANOG, Oct3/4, and Sox2 (primer sequences in Table S1). iPSCs also underwent karyotyping (Fig. S1) and whole-exome sequencing (WES). Reads were mapped to UCSC Genes hg19 and variants were called using GATK (software.broadinstitute.org/gatk/); variants were annotated using ANNOVAR (http://annovar.openbioinformatics.org/en/latest/). The capacity to differentiate into all three germ layers was examined *in vitro*. iPSCs were cultured on Matrigel-coated 12-well plates for three weeks in DMEM/F12 medium containing 20% KnockOut Serum Replacement, 1% GlutaMAX-I, 1% nonessential amino acids, 1X β-mercaptoethanol, and 1% penicillin-streptomycin. The differentiated cells were stained for markers of endoderm (AFP), mesoderm (SMA), and ectoderm (PAX6) (Fig. S2, antibodies in Table S2).

### CRISPR/Cas9-mediated *CHD8* KD

A CRISPR/Cas9 (clustered regularly interspaced short palindromic repeats/CRISPR associated protein 9) target of 20 nucleotides adjacent to the *CHD8* Ser62 codon (site of a nonsense mutation identified in an ASD patient) (21) was selected using the CRISPR Design website (crispr.mit.edu). This sequence for the single guide RNA was cloned into the pX330-U6-Chimeric_BB-CBh-hSpCas9 vector (Addgene plasmid #42230) (22). To facilitate homologous recombination, a donor DNA molecule was constructed as follows: the EGFP cassette (EGFP cDNA with upstream splice acceptor sequence, puromycin resistance gene, and CAG promoter) from the AAV-CAGGS-EGFP vector (Addgene plasmid #22212) (23) was amplified and inserted into the pCR2.1 cloning vector (Thermo Fisher Scientific) at the BamH1 and Not1 sites. 430 bp of upstream and 464 bp of downstream *CHD8* gene sequence and loxP sites were cloned to flank EGFP (Fig. 1a).

**Fig.1.**
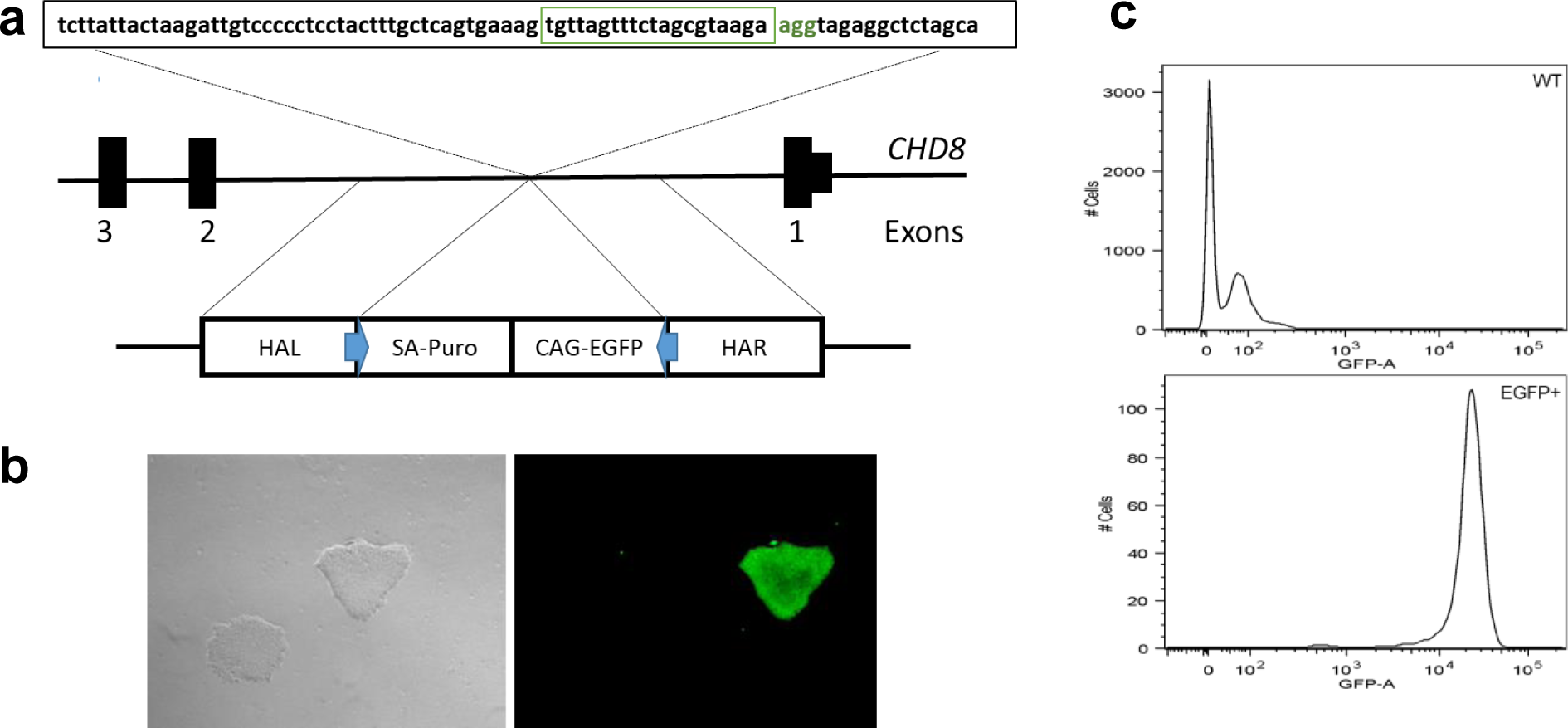
CRISPR/Cas9-mediated *CHD8* KD**. (a)** CRISPR/Cas9 target of 20 nucleotides (green box) and PAM site (green letters) adjacent to the *CHD8* Ser62 codon was used for the single guide RNA. To facilitate homologous recombination, a donor DNA molecule was used consisting of left (HAL) and right (HAL) homology arms and loxP sites (blue arrows) flanking the EGFP cassette (EGFP cDNA with upstream splice acceptor sequence (SA), puromycin resistance gene (Puro), and CAG promoter). **(b)** EGFP facilitated manual selection of successfully targeted iPSC colonies under fluorescence microscopy. **(c)** This was confirmed by fluorescence-activated cell sorting.

iPSCs were cultured in mTeSR1 in Matrigel-coated plates at 37°C, 5% CO2, and 90% humidity. At 70% confluency, iPSCs were harvested and dissociated into single cells using Accutase (STEMCELL Technologies). Using program A-023 of the Amaxa-4D Nucleofector (Lonza) and the Amaxa Human Stem Cell Nucleofector Kit 2 (VPH-5022, Lonza), one million cells were transfected with 2 μg CRISPR/Cas9 plasmid and 4 μg donor DNA plasmid. The transfected cells were plated in mTeSR1 and fed with fresh medium every other day. EGFP facilitated manual selection of successfully targeted iPSC colonies under fluorescence microscopy (Fig. 1b), which was confirmed by fluorescence-activated cell sorting (FACS) (Fig. 1c). Twelve EGFP+ colonies were split into two parts, one for expansion and one for DNA extraction and PCR amplification (primer sequences in Table S1). Homologous recombination of this cassette was expected to lead to premature truncation of CHD8. Each EGFP+ colony was found to have wild-type (WT) and mutant copies of *CHD8*.

Three separate WT and *CHD8* KD iPSC colonies were selected for further expansion. When cells from each of these colonies reached 70% confluence, they were digested by Accutase for seven minutes and repeatedly pipetted until a single cell suspension could be viewed under the microscope. FACS was used to sort and seed ∼1 EGFP+ cell per well of a 96-well plate. Medium was changed twice a week until cells were 60% confluent, which were then collected for DNA extraction and PCR amplification. Sanger sequencing confirmed the presence of WT and mutant copies of *CHD8*, which we expected to reflect largely heterozygous KD cells. An iPSC colony from one well for each of the three WT and *CHD8* KD iPSC lines was used for all subsequent studies. We sequenced the top 20 sites in the genome of the mutant iPSCs which were most similar to the CRISPR/Cas9 target region and confirmed the absence of off-target effects of genome editing at these sites (primer sequences in Table S1).

### Neural differentiation

iPSC clones were treated with 10 μM Y-27632 ROCK inhibitor (Tocris) and dissociated with Accutase. One million single cells were seeded per well of an AggreWell 800 plate (STEMCELL Technologies) and cultured for five days in neural induction medium (500 ng noggin, 10 μM SB43152, 1X N2, 1X B27, 200 mM l-glutamine, 1% penicillin-streptomycin). At day #5, embryoid bodies (EBs) were harvested and seeded in a 6-well plate coated with 15 μg/ml Poly-L-ornithine and 10 μg/ml Laminin (Sigma). The neural induction medium was changed every other day. When neural rosettes appeared, the medium was changed to 1X N2/B27 with 20 ng/ml FGF2 (Millipore). At 80-90% confluence, NPCs were dissociated with Accutase and maintained in 1X N2/B27 medium with 20 ng/ml FGF2 and 20 ng/ml EGF (PeproTech). For neural differentiation, NPCs were cultured in 1X N2/B27 medium supplied with 10 ng/ml BDNF, 10 ng/ml GDNF, 10 ng/ml IGF-1 (PeproTech) and 1 μM cAMP (Sigma). The medium was changed twice per week for 2-3 months to produce a mixed population of neuronal and glial cells (Fig. S2).

### Gene expression analysis

*RNA.* The level of *CHD8* mRNA KD was confirmed by qRT-PCR. Total RNA was isolated using the RNeasy kit (Qiagen), and purity and concentration were determined by the ND-1000 spectrophotometer (Nanodrop). cDNA was synthesized by the SuperScript First-Strand Synthesis System (Invitrogen). qRT-PCR was performed using SYBR Green (Roche) and the 2^−ΔΔCt^ method (24) (primer sequences in Table S1).

*Protein.* The level of CHD8 protein KD was measured by Western blot analysis. Whole-cell extracts from iPSCs or NPCs were obtained by treatment in lysis buffer (50 mM Tris pH 8.0, 140 mM NaCl, 1mM EDTA, 10% glycerol, 0.5% NP-40, 0.25% Triton X-100, 5mM DTT, 1mM PMSF, and protease inhibitor cocktail) followed by two minutes (min) of sonication. 15 μg of whole-cell extracts were mixed with Laemmli sample buffer containing 5% β-mercaptoethanol. Proteins were separated by 4-15% SDS-PAGE gel (Bio-Rad). The membranes were incubated with gentle shaking overnight at 4°C with rabbit anti-CHD8 (N-terminal) primary antibody (Cell Signaling Technology, Table S2) diluted 1:2000 in 5% (w/v) BSA, 1X TBS, and 0.1% Tween-20. They were then incubated for one hour at room temperature (RT) in HRP-conjugated donkey anti-rabbit secondary antibody (Jackson ImmunoResearch) diluted 1:10,0000. Actin was used as an internal control to measure the expression level of CHD8. HRP-conjugated anti-actin antibody (Cell Signaling Technology) was diluted 1:4000 in 5% (w/v) BSA, 1X TBS, and 0.1% Tween-20 and incubated overnight at 4°C. Labeled protein bands were visualized using SuperSignal West Pico Chemiluminescent Substrate (Thermo Fisher Scientific) under the ChemoDoc XRS system, and quantitated using Bio-Rad Image Lab 6.0.

### Immunocytochemistry and neuronal morphology

Cells were fixed in 4% paraformaldehyde at RT for 10 min and then permeabilized with 0.1% Triton X-100 in PBS for 15 min. They were blocked in 5% goat serum for one hour before incubation with primary antibody (in 1X PBS, 1% BSA, 0.1% Triton X-100) at 4°C overnight. After 3 washes in PBS/0.1% Triton X-100, cells were incubated with secondary antibody (in 1X PBS, 1% BSA, 0.1% Triton X-100) at RT for one hour (antibodies in Table S2). Fluorescent signals were visualized under an inverted microscope (Leica), and images were processed using ImageJ (imagej.nih.gov/ij). Signals for microtubule-associated protein 2 (MAP2), glial fibrillary acidic protein (GFAP), Synapsin 1 (SYN1), and vesicular glutamate transporter 1 (VGLUT1) were counted by Puncta Analyzer (25). The Neurite Outgrowth Staining Kit (Invitrogen) was used to label live neurites. Cells were incubated with 50 μl of a solution containing 0.1 μg/ml Hoechst dye and 1:1000 cell membrane stain for 30 min. After the staining solution was removed, 50 μl of background suppression dye was applied. Images were taken under the inverted microscope; neurite length and root number were measured by ImageJ.

### Proliferation assay

The effect of CHD8 KD on NPC proliferation was examined using the Cell Proliferation Kit I (MTT) (Roche). Cells were seeded in a 96-well plate at a density of 8×10^3^ cells/well. After one, two, and three days in culture (37°C, 5% CO_2_, 90% humidity), 50 μg MTT was added to each well and incubated for four hours. 100 μl solubilization solution was added to each well and incubated overnight. Absorbance was measured at 570 nm with a reference reading at 750 nm using the GloMax-Multi Detection System (Promega).

### Cell cycle analysis

The cell cycle of NPCs was analyzed by using the Click-iT Plus EdU Alexa Fluor 647 Flow Cytometry Assay Kit (Thermo Fisher Scientific). 5 uM EdU was added to one million cells in a 6-well plate for four hours. The medium was changed to remove unabsorbed EdU, and incubation was continued overnight (37°C, 5% CO2, 90% humidity). Cells were then subjected to FACS, and the data were analyzed using FlowJo software (flowjo.com).

### Statistics

All data was obtained from three biological replicates for each iPSCs, NPCs, and neural cells. All assays were performed on three technical replicates of each sample. Data are expressed as mean ± standard deviation (SD). Comparisons were performed using the t-test; *P* < 0.05 was considered statistically significant.

## RESULTS

### Characterization of *CHD8* KD cell lines

Approximately three weeks after activated WT human T-lymphocytes were infected with Sendai virus encoding cMyc, Klf4, Oct3/4, and Sox2, iPSC colonies were isolated for passaging and characterization. Pluripotency was confirmed by staining for ALP, Tra-1-60, SSEA3, and SSEA4 (Fig. S2). RT-PCR showed that the iPSCs had active endogenous expression of cMyc, Klf4, NANOG, Oct3/4, and Sox2 but no genomic incorporation of the virally delivered factors. Loss of Sendai virus sequences was confirmed by PCR (Table S1). The iPSCs showed normal karyotypes (Fig. S1), and WES showed no rare *CHD8* coding variants. They showed capacity for differentiating into all three germ layers *in vitro* by staining positively for the markers AFP (endoderm), SMA (mesoderm), and PAX6 (ectoderm) (Fig. S2).

The ASD patient S62X mutation is located in the first exon of *CHD8* (21). We modeled this mutation in iPSCs by using CRISPR/Cas9 genome editing and homologous recombination to insert a cassette in the first intron to yield premature truncation of the CHD8 protein and expression of the *EGFP* gene (Fig. 1). Sanger sequencing showed that each EGFP+ iPSC colony contained both WT and mutant copies of *CHD8*. Three separate WT and *CHD8* KD iPSC colonies were subsequently differentiated into NPCs and neural cells (mixed population of neuronal and glial cells, Fig. S2). qRT-PCR showed that, compared to WT, *CHD8* mRNA levels were reduced in KD cell lines by an average of 41% in iPSCs, 52% in NPCs, and 59% in neural cells (Fig. 2a). Western blot analysis of NPCs showed that CHD8 protein was reduced by an average of 58% (Fig. 2b and 2c).

**Fig. 2.**
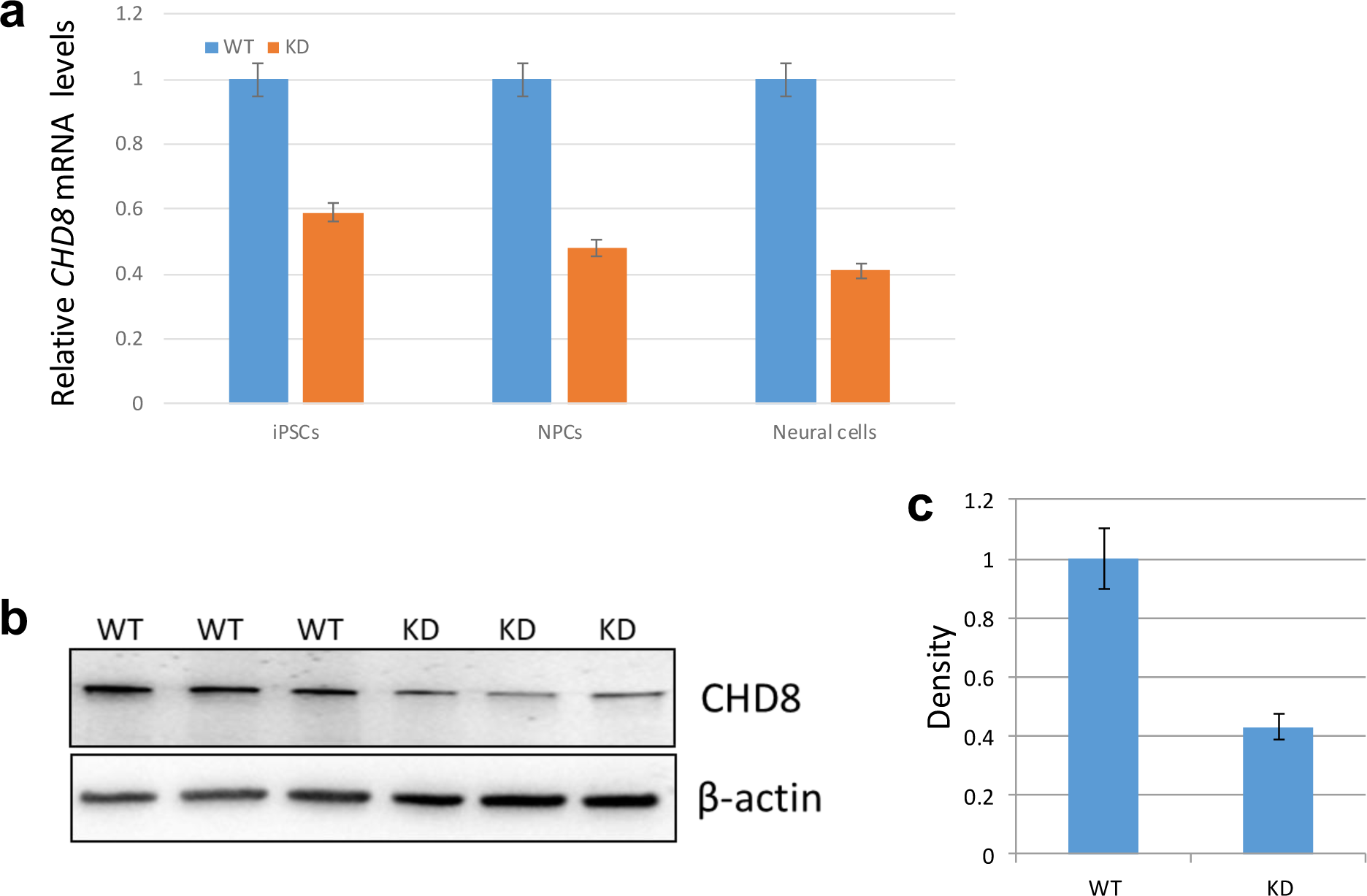
Gene expression analysis. **(a)** qRT-PCR showed that, compared to WT, *CHD8* mRNA levels were reduced in KD cell lines by an average of 41% in iPSCs, 52% in NPCs, and 59% in neural cells. **(b and c)** Western blot analysis of NPCs showed that CHD8 protein was reduced by an average of 58%.

### Effects of *CHD8* KD on cell proliferation and differentiation

During the colony formation of iPSCs, we noted that *CHD8* KD led to an increased number of iPSC colonies. Two weeks after 50,000 WT and *CHD8* KD iPSCs were plated, an average of three colonies were generated by WT iPSCs while an average of 20 colonies were generated by the mutant iPSCs (*P* = 0.00231, t-test, Fig. 3a). After colony formation, it is typical to see areas of spontaneous differentiation along the edges of colonies during routine maintenance. Two weeks after colony formation, we observed fewer areas of differentiation around mutant iPSCs compared to WT iPSCs (Fig. 3c). We measured the proliferation of NPCs by the MTT assay and found a significant increase in proliferation of *CHD8* KD NPCs (0.473 ± 0.040, day 4) compared to WT NPCs (0.292 ± 0.019, day 4) (*P* = 3.34 × 10^−5^, t-test, Fig. 3b).

**Fig. 3.**
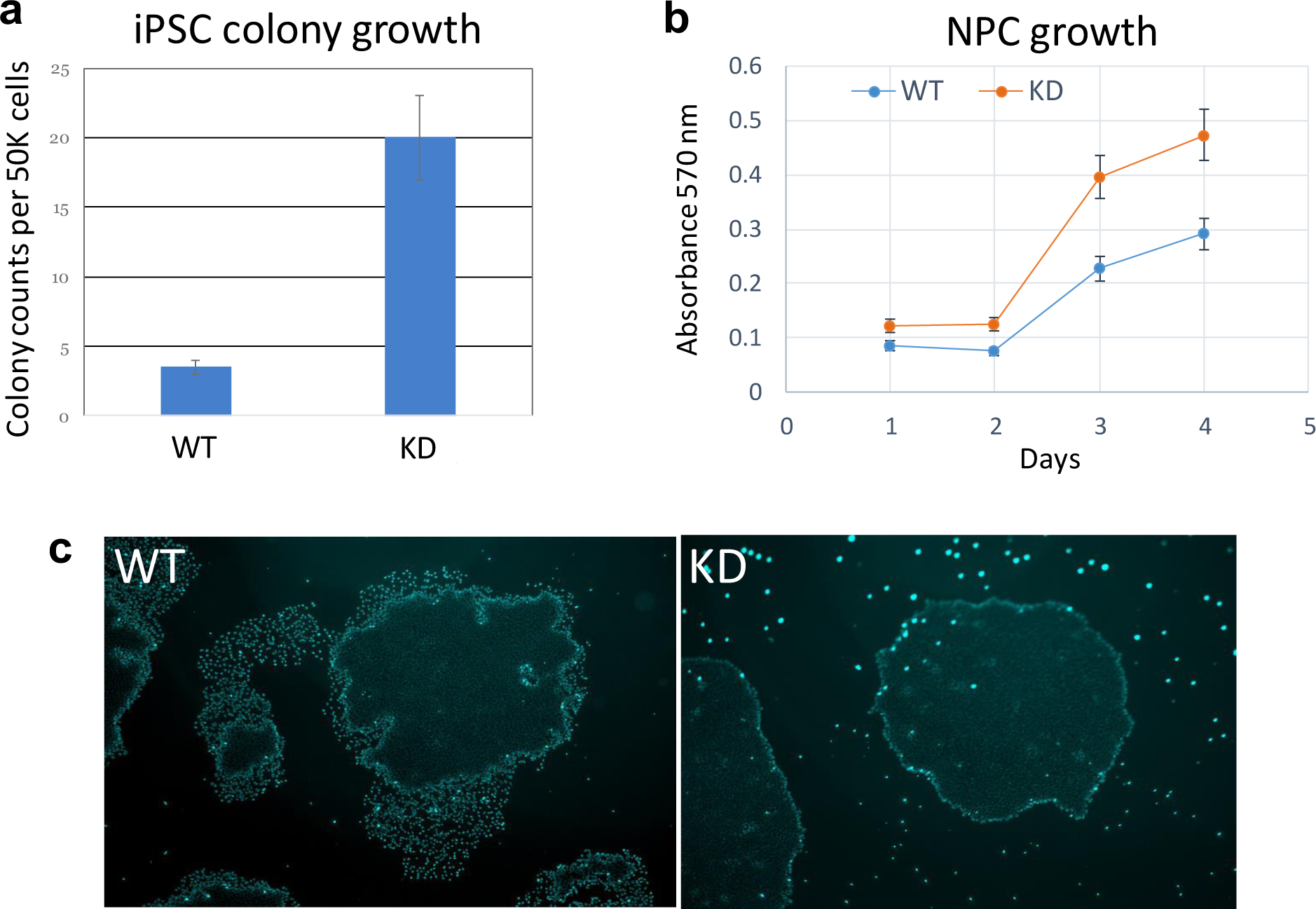
Effects of *CHD8* KD on proliferation and differentiation. *CHD8* KD caused **(a)** increased iPSC colony numbers, **(b)** increased NPC growth, and **(c)** suppression of spontaneous differentiation around the edges of iPSC colonies.

During the differentiation of iPSCs to NPCs, cells typically organize into rosettes by approximately two weeks. We observed that the formation of rosettes was delayed for *CHD8* KD cells compared to WT cells by approximately one week (Fig. 4). Three days after WT NPCs were plated, we noted the outgrowth of neurites. By 12 days, neurites covered the whole field of observation. For *CHD8* KD NPCs, we did not observe the outgrowth of neurites until six days after plating, and on day #12, only partial sprouting was seen. We calculated a significant decrease in neurite area per mutant cell (nuclear) area (138.23 ± 2.54) compared to WT cell (nuclear) area (355.56 ± 31.62, *P* = 0.025, t-test, Fig. 5). Given these observations on the effects of *CHD8* KD on cell proliferation and differentiation, we performed cell cycle analysis of NPCs. A significantly smaller percentage of *CHD8* KD NPCs (58.84 ± 4.59) is found in the G0/G1 phase of the cell cycle compared to WT NPCs (72.16 ± 6.8, *P* = 0.007, t-test), and a larger percentage of *CHD8* KD NPCs (21.90 ± 7.08) is found in the G2/M phases of the cell cycle compared to WT NPCs (12.10 ± 6.82, *P* = 0.057, t-test, Fig. 6).

**Fig. 4.**
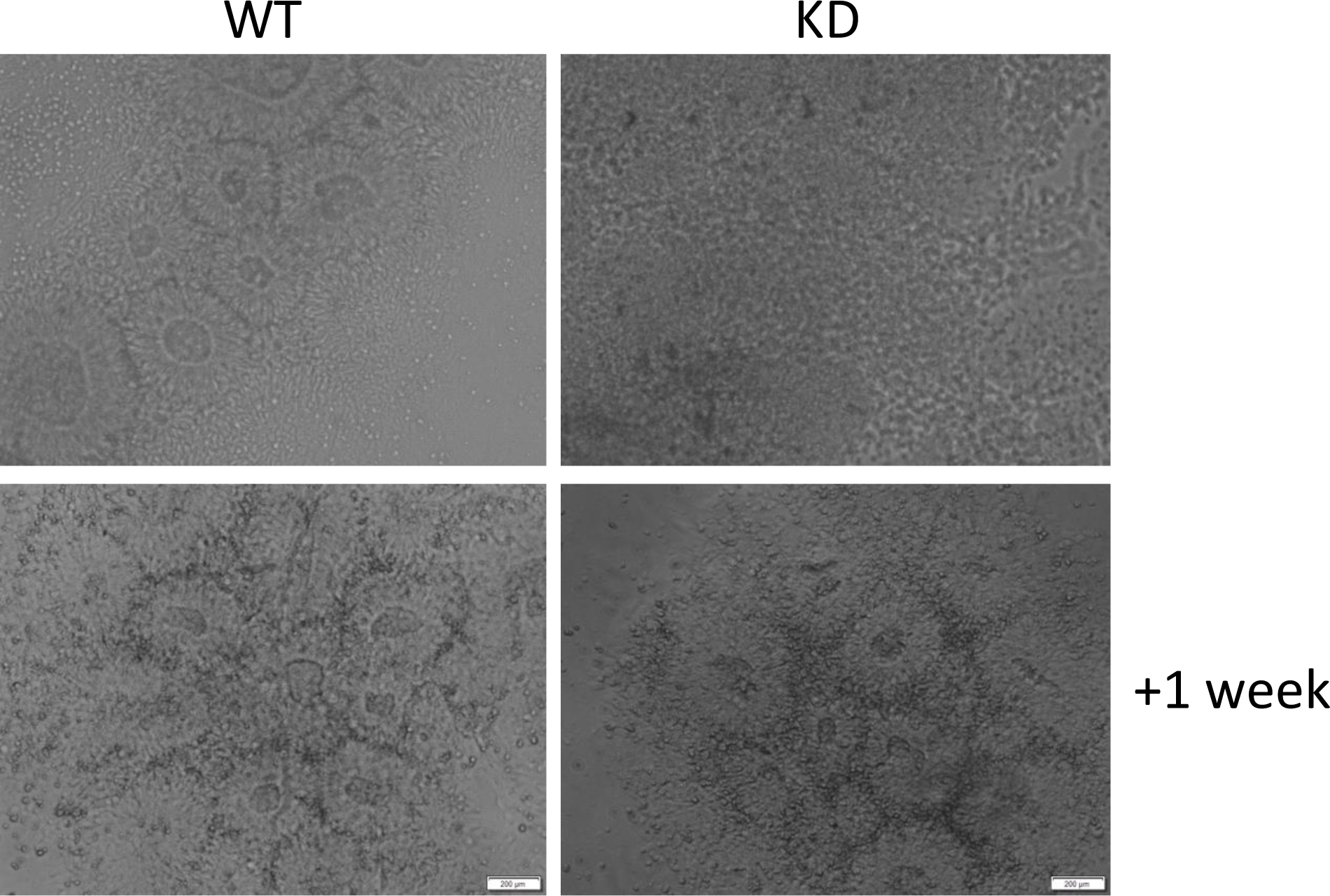
NPC rosette formation. During the differentiation of iPSCs to NPCs, cells typically organize into rosettes by approximately two weeks. *CHD8* KD delayed rosette formation by one week, or by approximately a third.

**Fig. 5.**
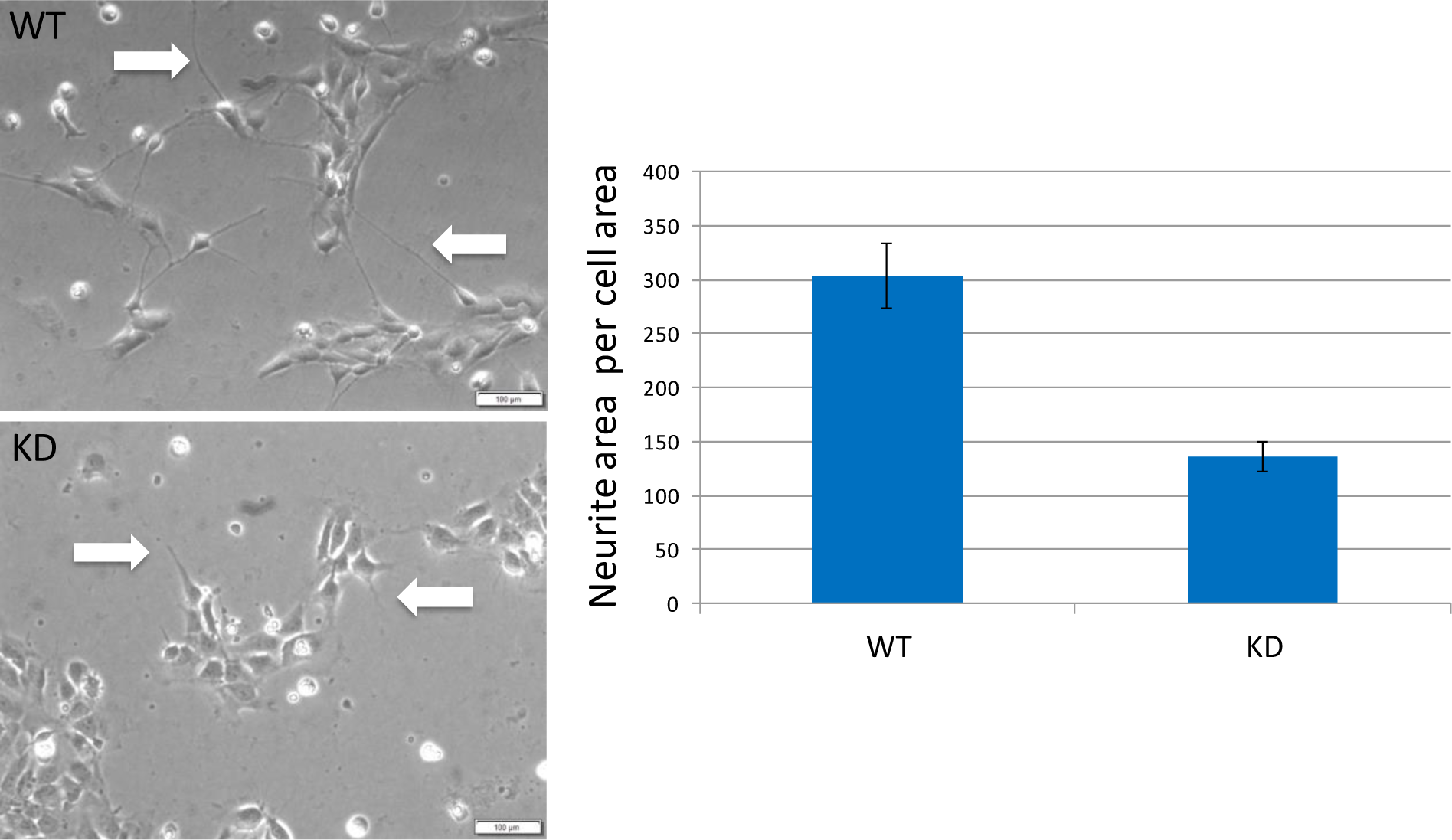
Neurite outgrowth. 12 days after NPCs were plated, *CHD8* KD resulted in a significantly decreased neurite area (white arrows) per cell (nuclear) area.

**Fig. 6.**
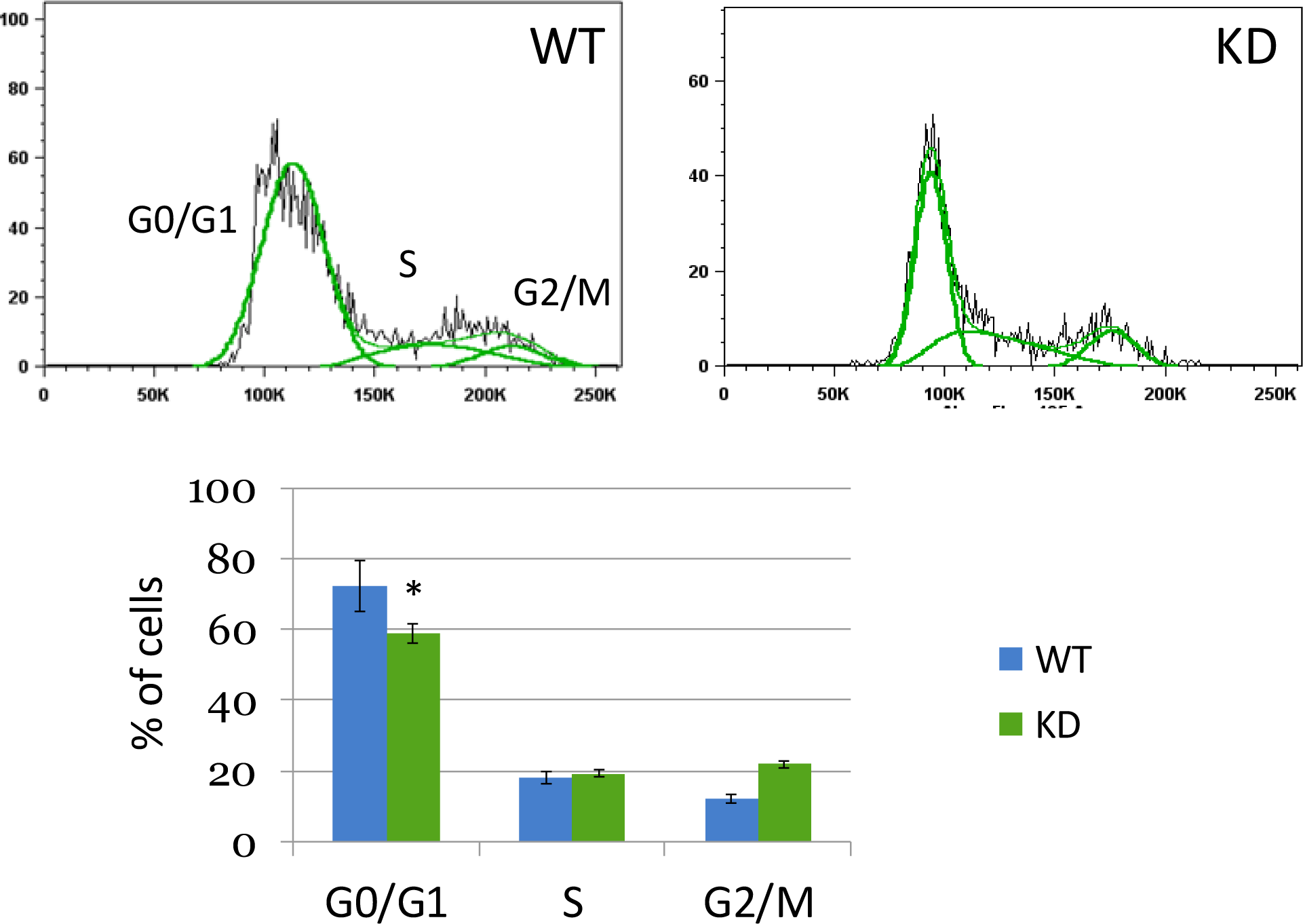
Effect of *CHD8* KD on the cell cycle. A significantly smaller percentage of *CHD8* KD NPCs (58.84 ± 4.59) is found in the G0/G1 phase of the cell cycle compared to WT NPCs (72.16 ± 6.8, *P* = 0.007, t-test), and a larger percentage of *CHD8* KD NPCs (21.90 ± 7.08) is found in the G2/M phases of the cell cycle compared to WT NPCs (12.10 ± 6.82, *P* = 0.057, t-test).

### Effects of *CHD8* KD on neural cell composition

Fifty days after NPCs were differentiated into a mixed population of neuronal and glial cells, we investigated the composition of this mixed population by immunocytochemistry. First, we determined the ratio of neurons to glial cells by staining for the neuronal marker MAP2 and the glial marker GFAP. WT neural cells (6.55 ± 1.20) had a significantly greater ratio of MAP2 to GFAP staining than the *CHD8* KD cells (2.60 ± 0.50, *P* = 0.0001, t-test, Fig. 7a). This increase in ratio (6.55/2.60 = 2.52x) is due to a similar increase in MAP2 staining in WT versus *CHD8* KD neurons (2.89x) rather than a difference in GFAP staining (1.15x). Next, we determined the fraction of MAP2+ cells which were also positive for the excitatory neuronal marker VGLUT1. A significantly greater fraction of WT MAP2+ cells stained for VGLUT1 (0.68 ± 0.19) than of *CHD8* KD MAP2+ cells (0.44 ± 0.12, *P* = 0.003, t-test, Fig. 7b). Finally, we examined the presence of the synaptic marker SYN1 among the neuronal cells. A significantly greater percentage of WT MAP2+ cells also stained for SYN1 (7.55 ± 0.80) than of *CHD8* KD MAP2+ cells (1.89 ± 0.10, *P* = 0.029, t-test, Fig. 7c).

**Fig. 7.**
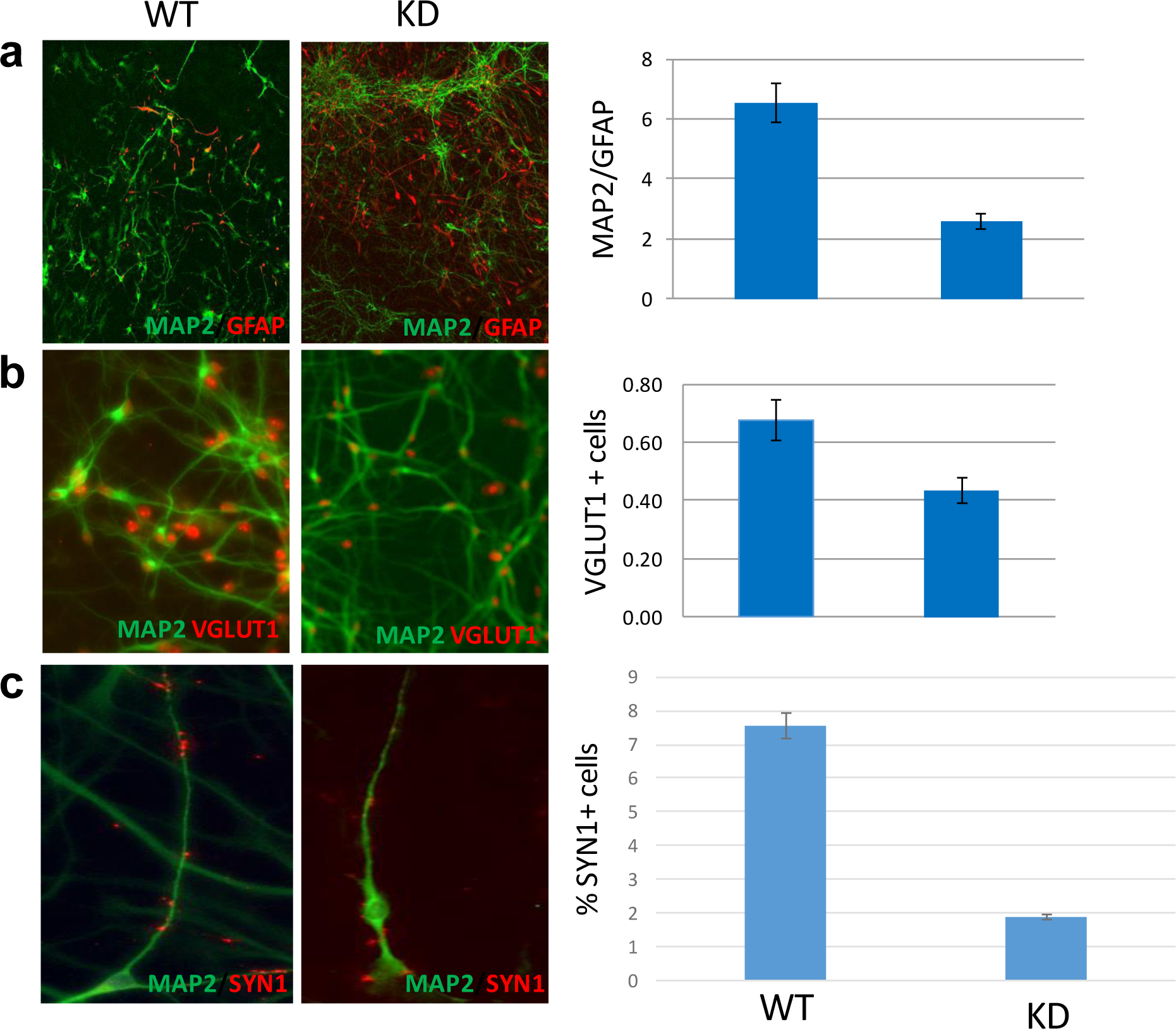
Effects of CHD8 KD on cell composition. CHD KD resulted in: **(a)** decreased MAP2/GFAP ratio, **(b)** decreased fraction of VGLUT1+ cells, and **(c)** decreased percentage of SYN1+ cells.

## DISCUSSION

*CHD8*, which encodes a chromatin remodeling protein that regulates Wnt/β-catenin mediated gene expression (6, 7), is one of the most strongly associated genes with ASD (5). A common emerging theme of *CHD8* disruption across ASD patients and animal models is macrocephaly (5). Macrocephaly may be due to different factors, such as over proliferation of neural cells or increased neural cell body size. Our goal was to generate a human stem model of *CHD8* deficiency and investigate basic properties of neural proliferation and differentiation to provide insight into how patient mutations contribute to the pathophysiology of ASD at the cellular level.

We made a series of observations which implicate *CHD8* deficiency in alterations of the cell cycle. More specifically, *CHD8* KD increased cell proliferation and delayed neural differentiation by: (1) increasing the number of iPSC colonies formed, (2) suppressing spontaneous differentiation along the edges of iPSC colonies, (3) increasing the proliferation of NPCs, (4) delaying the formation of neural rosettes, (5) delaying neurite outgrowth, (6) decreasing the percentage of cells in the G0/G1 phase of the cell cycle, and (7) increasing the percentage of cells in the G2/M phase of the cell cycle. The effects of *CHD8* deficiency on neural cell composition reflected delayed neural differentiation as *CHD8* KD resulted in: (1) decreased presence of the neuronal marker MAP2 although not the glial marker GFAP, (2) decreased presence of the excitatory neuronal marker VGLUT1, and (3) decreased presence of the synaptic marker SYN1. It should be noted, however, that the role of CHD8 in regulating the cell cycle is likely more complicated than suggested by our study. In contrast to our results, CHD8 depletion in RPE1 cells caused a delay in cell cycle progression (26). Furthermore, both *CHD8* overexpression and deficiency have been detected in various cancers (5). Differences in the role of *CHD8* may be due to the specific cell type.

While increased cell proliferation may underlie the macrocephaly seen in ASD patients and animal models, we cannot rule out an effect of *CHD8* KD on increasing neural cell body size, which can be determined by future, more detailed studies of neural morphology. Delayed neural differentiation may also contribute to the pathophysiology of ASD. The observation that *CHD8* KD decreased the number of VGLUT1+ cells is interesting in light of the theory that there is an imbalance in excitation/inhibition (E/I) in ASD. Initially, it was proposed that an increased ratio of E/I may be caused by a combination of genetic and environmental factors that disrupt key neural systems in ASD (27). However, a recent study of four diverse ASD mouse models suggests that increased E/I reflects a compensatory change rather than causing the disorder (28). It will be important to investigate this possibility in the various mouse models of *CHD8* deficiency that have been generated. We note that we were unable to detect inhibitory markers in 50 day-old neuronal cells to investigate the effect of *CHD8* KD on this cell population; this will require extended culturing in future studies.

A limitation of our study is that, although *CHD8* mRNA and protein levels were reduced by approximately half in the KD lines, we cannot rule out that each of our iPSC colonies were mosaics of homozygous WT and heterozygous KO cells. It was technically challenging to create pure clones since iPSCs do not survive single cell passaging well in our experience. A study by an independent group was not able to generate homozygous *CHD8* KO in human iPSCs, suggesting that lack of CHD8 is lethal (20), as it is in mice (10). Therefore, it is unlikely that our colonies contained homozygous *CHD8* KO cells, which could theoretically cause our cellular phenotypes to be more abnormal than those of heterozygous KO cells, which we aimed to model as a reflection of the heterozygous mutations identified in ASD patients. Instead, the presence of WT cells may have caused us to underestimate the abnormality of the cellular phenotypes we observed.

In summary, we generated and compared WT and *CHD8* KD human iPSCs, NPCs, and neural cells. Our results suggest that CHD8 deficiency causes increased cell proliferation and delayed neural differentiation, which may contribute to the pathophysiology of ASD. In addition to the future studies mentioned above, an important next step will be to investigate the electrophysiological properties of the neural cells. Also, the introduction of a *CHD8* mutation in WT human iPSCs demonstrated that CHD8 deficiency is sufficient to cause cellular abnormalities. The generation of iPSCs and neural cell lines from an ASD patient with a *CHD8* mutation and correction by CRISPR/Cas9 genome editing will test the hypothesis that *CHD8* disruption is necessary to cause cellular abnormalities, strengthening the evidence for the association of *CHD8* with ASD.

## ADDITIONAL FILES

**Additional file 1: Fig. S1.** Human T-lymphocytes, iPSCs, and karyotype

**Additional file 2: Fig. S2.** Generation and immunostaining of human iPSCs, NPCs, and neural cells

**Additional file 3: Table S1.** Primers sequences

**Additional file 4: Table S2.** Antibodies

## ABBREVIATIONS

ASD: Autism spectrum disorder
ALP: Alkaline phosphatase
CHD8: Chromodomain helicase DNA-binding protein 8
CRISPR/Cas9: Clustered regularly interspaced short palindromic repeats/CRISPR associated protein 9
EB: Embryoid body
FACS: Fluorescence-activated cell sorting
GFAP: Glial fibrillary acidic protein
GI: Gastrointestinal
iPSC: Induced pluripotent stem cell
KD: Knockdown
KO: Knockout
MAP2: Microtubule-associated protein 2
MEF: Mouse embryonic fibroblast
NPC: Neural progenitor cell
PBMC: Peripheral blood mononuclear cell
SYN1: Synapsin 1
VGLUT1: Vesicular glutamate transporter 1
WES: Whole-exome sequencing
WT: Wild-type

## ACKNOWLEDGEMENTS

We are thankful to our healthy male volunteer for the blood sample to generate iPSCs and to our colleagues at the Yale Stem Cell Center for guidance: Yinghong Ma, Caihong Qiu, and Jason Thomson.

## FUNDING

This research was funded by the Connecticut Regenerative Medicine Research Fund (15-RMA-YALE-32 to ARG).

### Availability of data and materials

The data generated and analyzed during this study are included in this article (and its additional files) and are available from the corresponding author upon request.

### Authors’ contributions

WL designed the study, generated and characterized all cell lines, and performed bioinformatics analysis. WD performed bioinformatics analysis. EJH and TVF analyzed data. ARG conceived and designed the study, analyzed the data, and wrote the manuscript. All authors read, edited, and approved the final manuscript.

### Consent for publication

The IRB protocol provides for consent for publication. However, no individually identifying information is presented in this report.

### Ethics approval and consent to participate

This research was approved by the Yale University IRB, and informed consent was obtained from the subject (Human Investigations Committee (HIC) Protocol # 0301024156).

---

**Fig. S1.**
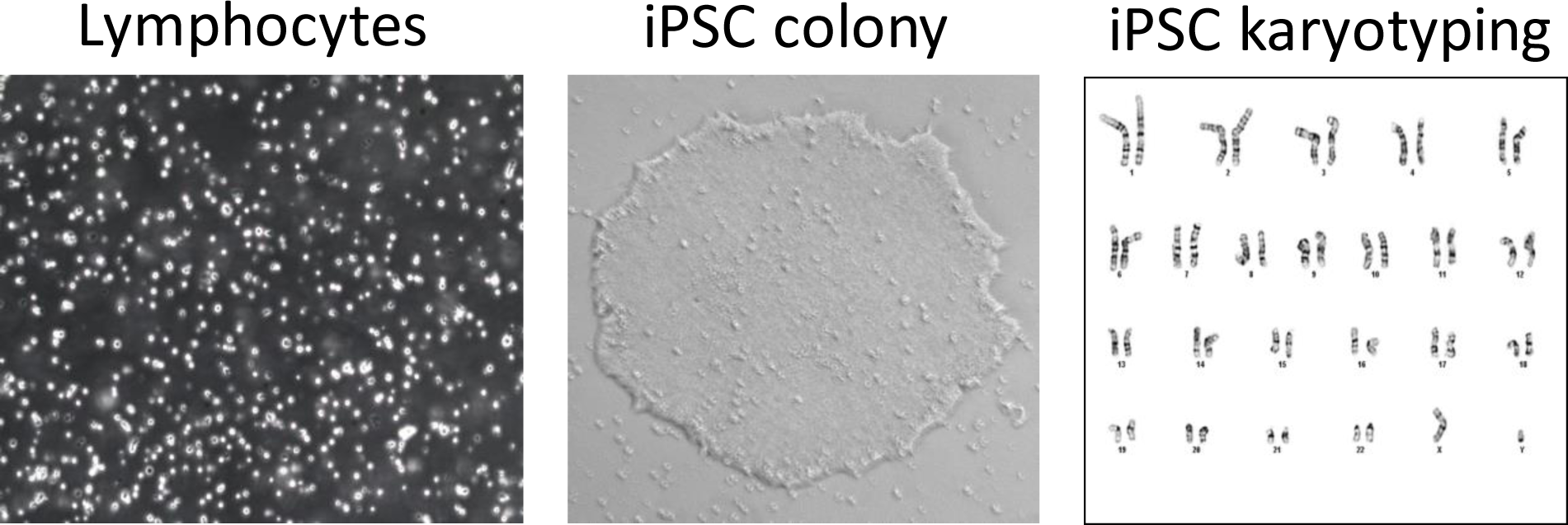
Human T-lymphocytes were reprogrammed into iPSCs. WT and *CHD8* KD karyotypes were normal.

**Fig. S2.**
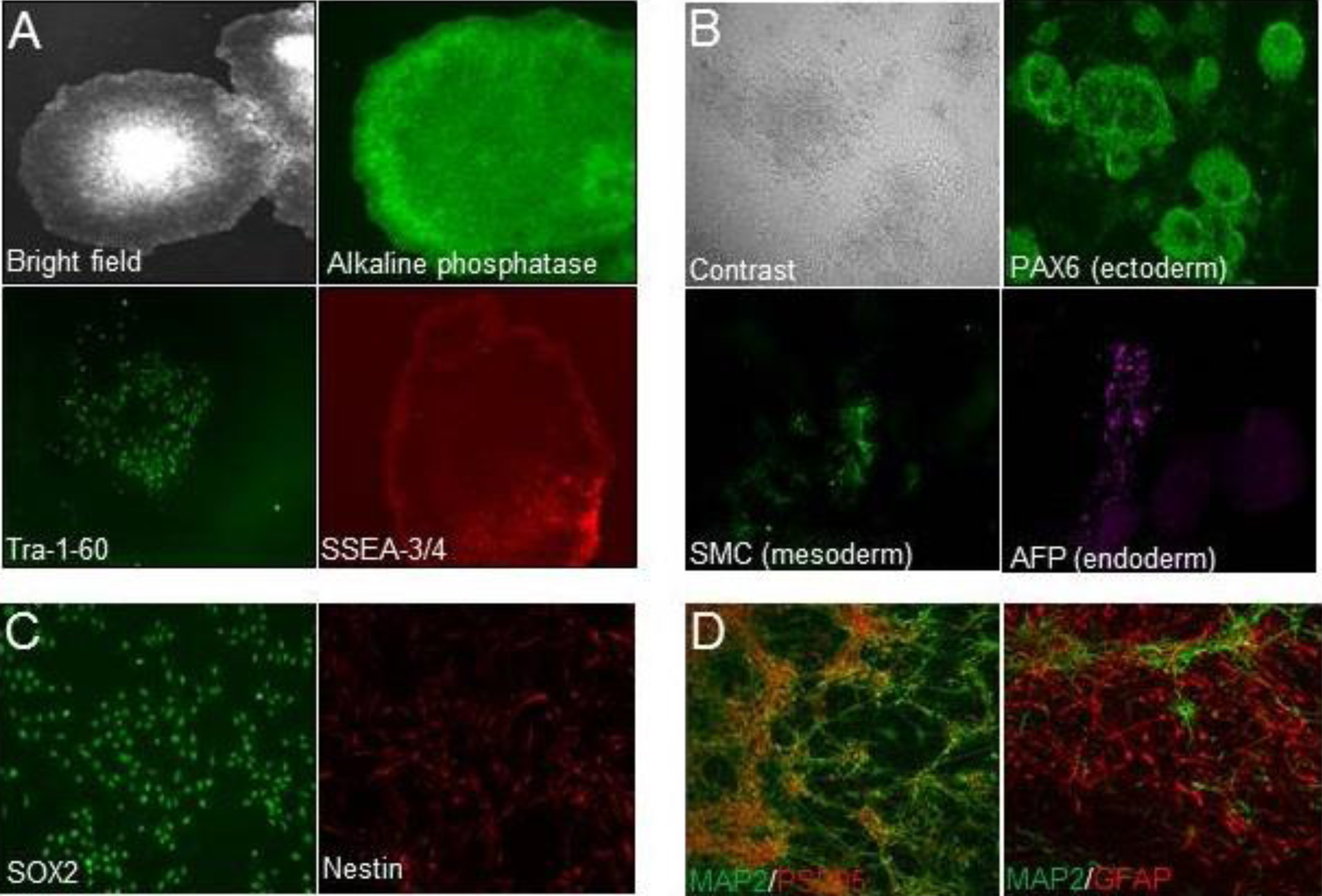
Creation of cell lines. Characterization of iPSCs by **(A)** staining for surface markers and **(B)** formation of all three germ layers. The iPSCs were differentiated into **(C)** NPCs. The NPCs were, in turn, differentiated into **(D)** neural cells.

